# Lineage-dependent differences and the role of IFITM3 in the type-I interferon-induced restriction of Zika virus

**DOI:** 10.1101/455972

**Authors:** Theodore Gobillot, Daryl Humes, Amit Sharma, Julie Overbaugh

**Affiliations:** Division of Human Biology, Fred Hutchinson Cancer Research Center, Seattle, Washington, United States of America; Medical Scientist Training Program, University of Washington School of Medicine, Seattle, Washington, United States of America

## Abstract

Type-I interferon (IFN-I) is an important aspect of host innate antiviral response. Recent studies have shown that IFN-I can inhibit Zika virus (ZIKV) replication and that this is mediated in part by Interferon-induced transmembrane protein 3 (IFITM3). ZIKV infections in South America have led to severe congenital syndrome in a subset of infected infants. ZIKV was first identified in Africa, where there is limited evidence for the pathogenic effects associated with the American outbreak, which is fueled by infection with Asian-lineage strains, raising the possibility that the African and Asian ZIKV lineages have distinct pathogenic properties. Given the observation that IFN-I can inhibit ZIKV replication in cell culture, we asked whether ZIKV strains differed in their susceptibility to IFN-I. There was a range of susceptibilities to IFN-I inhibition across virus strains. Virus production in A549 cells was reduced from 3-42-fold for IFNα and 63-807-fold for IFNβ across a panel of nine viruses, five from the African-lineage and four from the Asian-lineage. African-lineage ZIKV strains were more resistant to IFN-I than Asian-lineage strains, but this difference was only significant for IFNα-mediated restriction (p = 0.049). Notably, over-expression of IFITM3 at similar levels induced by IFN-I did not significantly restrict either a prototype African lineage (MR 766) or Asian lineage (PRVABC59) isolate. Moreover, knocking out IFITM3 expression did not result in a significant increase in viral replication or a diminishment of the inhibition by IFN-I. Overall, our findings show that while diverse ZIKV strains are susceptible to the antiviral effects of IFN-I, African-lineage strains are more resistant to IFNα. In addition, the majority of the IFN-I-induced inhibition of ZIKV strains cannot be explained by IFITM3, suggesting that other unknown ISGs may be the driving force of the type I IFN response against ZIKV.

**Author summary:** The innate immune system, and specifically the type-I interferon response, is a critical component of the host response against viral infections. The recent unprecedented spread and severe pathogenic features of Zika virus in the Americas have led to significant interest in characterizing features of Zika virus strains that have fueled the American outbreak. Zika virus was first identified in Africa, where there is limited evidence for the pathogenic effects associated with the American outbreak. Here, we demonstrate that African-lineage Zika virus strains are significantly more resistant to the effects of type-I interferon, and that type-I interferon-mediated restriction of Zika virus strains is not explained by the host factor Interferon-induced transmembrane protein 3. This improved understanding of Zika virus-host interactions may explain certain pathogenic features of Asian-lineage Zika virus strains that have fueled the American Zika virus epidemic, and supports the search for as-yet-unidentified actors in the interferon response against Zika virus.

## Introduction

The recent spread and severe pathogenic features of Zika virus (ZIKV) in the Americas have highlighted the epidemic potential of this emerging pathogen. ZIKV was detected in Brazil in May 2015 [1]. By December 2015, ZIKV had infected an estimated 1.3 million individuals in the region [2]. During this time, an American outbreak clade of ZIKV strains clustered within the Asian-lineage was linked to fetal abnormalities, a severe congenital syndrome in neonates, and adverse neurological outcomes in adults [3-6]. Prior to the American epidemic, only two outbreaks of ZIKV had been reported. In 2007, ZIKV first emerged in the Pacific on Yap Island and infected ~75% of the island’s population [7]. From 2013 to 2014, an outbreak of ~30,000 symptomatic ZIKV infections was reported in French Polynesia, which then rapidly spread to other Pacific Islands [8, 9].

ZIKV was first identified in Africa over 70 years ago and only sporadic infections were reported from tropical Africa and Asia prior to its emergence in the Pacific. There is limited evidence that African-lineage ZIKV infections are associated with the severe pathogenic profile that has been described in recent ZIKV outbreaks fueled by Asian-lineage strains. This raises the possibility that African- and Asian-lineage ZIKV strains may have distinct pathogenic properties. Interestingly, a number of studies have suggested that African lineage strains tend to have increased replication kinetics, cytopathicity and more severe pathogenic outcomes in small animal models as compared to Asian-lineage strains [10-22]. This increased replication fitness of African lineage viruses is somewhat surprising given the relative absence of disease associated with this viral clade, raising additional questions about mechanisms of ZIKV pathogenesis.

Type-I interferon (IFN-I) is a critical component of the host innate immune response to viral infection [23]. Upon recognition of viral infection, target cells enter a transcriptional program that increases the production of IFN-I (IFNα and IFNβ), which establishes an anti-viral state in bystander cells and restricts viral replication in infected target cells [24]. The ability of IFN-I to restrict viral replication is largely due to the activation of thousands of interferon-stimulated genes (ISGs) that have a wide range of anti-viral functions [25]. IFN-I is capable of restricting ZIKV in cell culture [26, 27], and most murine models of ZIKV infection and pathogenesis require ablation of the IFN-I signaling pathway, underscoring the important role of ISGs in restricting ZIKV replication [19]. One such ISG is the Interferon-Induced Transmembrane Protein 3 (IFITM3), which was the first ISG described as a key effector of the IFN-I response against ZIKV [28, 29]. IFITM3 is a small transmembrane protein that restricts a broad array of viruses and is potently induced by IFN-I [30]. It is unclear whether strains of ZIKV differ in their susceptibility to IFN-I-mediated restriction.

The goals of this study were to determine whether Zika viruses differ their susceptibility to restriction by IFN-I and whether there are overall differences between African and Asian lineage viruses. We demonstrate that African-lineage viruses are significantly more resistant to the effects of IFNα than Asian-lineage viruses; they also show greater resistance to IFNβ, but the difference is not significant as strains from both lineages are potently restricted by IFNβ. We also find that IFITM3 does not explain the IFN-I-mediated restriction of the nine ZIKV strains tested. These findings support the continued identification and characterization of additional IFN-I-induced factors that restrict ZIKV.

## Results

### Effect of IFN-I treatment on diverse ZIKV strains in A549 cells

In order to test the hypothesis that IFN-I sensitivity differs between African-lineage and Asian-lineage ZIKV strains, a panel of nine viruses was tested for their ability to replicate in A549 cells in the presence or absence of IFN-I. Five strains (MR 766, IbH 30656, DAK-AR-25, DAK-AR-67, DAK-AR-71) belong to the African lineage and four strains (FLR, PRVABC59, H/PAN/2016/BEI, H/PAN/2015/CDC) belong to the American outbreak clade within the Asian lineage (Fig 1, blue asterisks). The percent identity of the complete genomes of African vs. Asian lineage strains in this panel is 88-89%, which is representative of the overall diversity of isolated ZIKV strains [8]. All Asian-lineage viruses were isolated from infected humans, while only one African-linage virus was isolated from an infected human (IbH 30656). Three African-lineage strains were isolated from mosquitoes (DAK-AR-25, DAK-AR-67, DAK-AR-71) and one from a sentinel rhesus macaque (MR 766) (Table 1). In addition, these strains have diverse passage histories. Most have undergone 3-5 passages in mosquito (AP61, C6/36) and/or African-green monkey (Vero) cell lines; however, MR 766 has been extensively passaged in mouse brain and subsequently in Vero cells. IbH 30656 has a similar but less extensive high-passage profile. Of note, the number of passages in AP61 cells in DAK-AR-67 and DAK-AR-71 is unknown.

**Fig 1.**
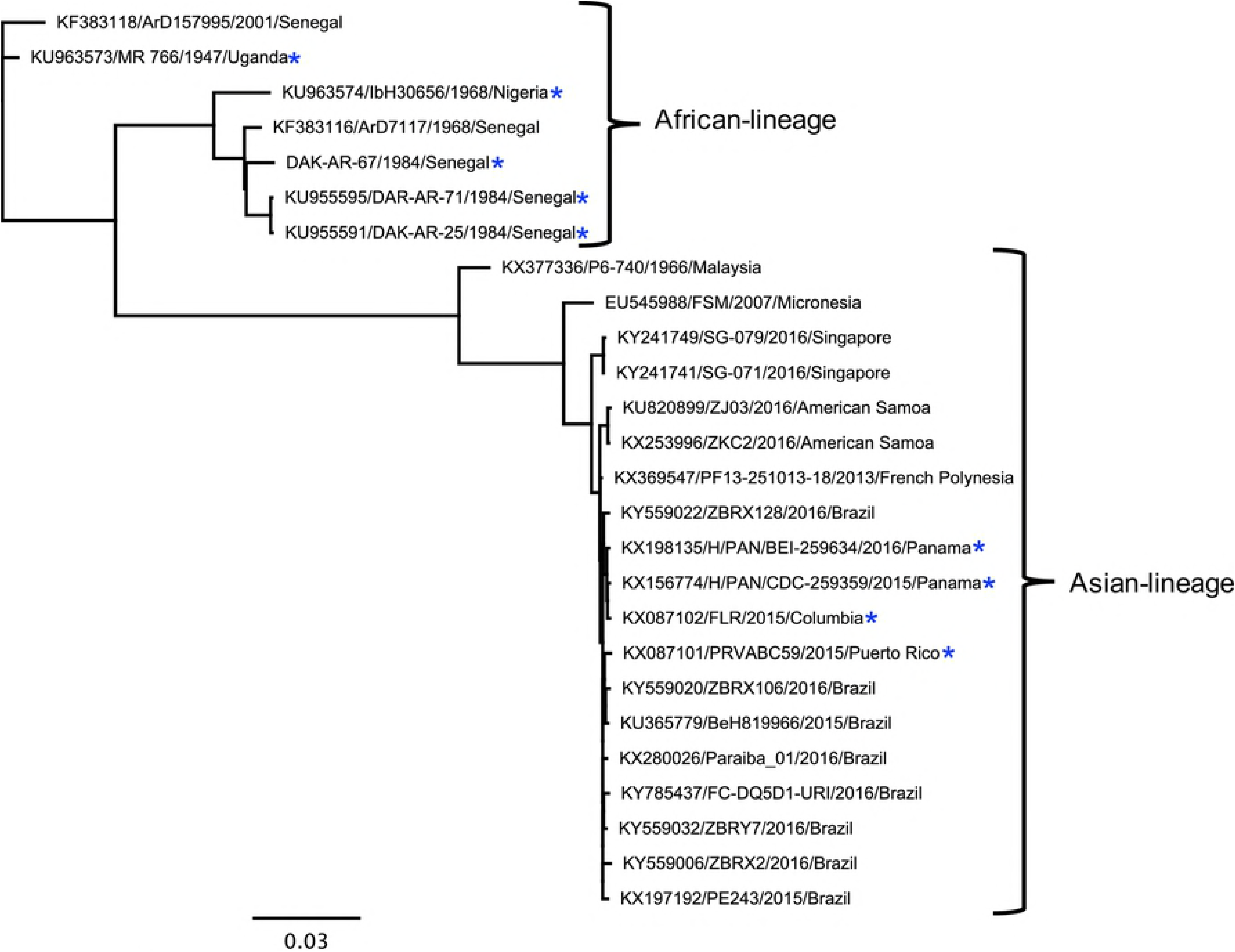
Phylogenetic relationships of Zika virus strains used in this study. Maximum-likelihood phylogeny of full-length open-reading-frame nucleotide sequences using Zika virus strains in this study (blue asterisks) and reference sequences isolated from humans, non-human primates, and mosquitoes. At least one representative strain from each documented ZIKV clade is included in the phylogenetic tree.

**Table 1.** Summary of characteristics of ZIKVs used in this study.

| *Strain* | *Lineage* | *Source* | *Passage History* |
| --- | --- | --- | --- |
| MR 766 | African | Rhesus (Uganda 1947) | 150x mouse brain<br>8x Vero cells |
| IbH 30656 |  | Human (Nigeria 1968) | 21x mouse brain<br>4x Vero cells |
| DAK-AR-25 |  | *Aedes africanus* (Senegal 1984) | 1x AP61 cells<br>1x C6/36 cells<br>3x Vero cells |
| DAK-AR-67 |  | *Aedes taylori* (Senegal 1984) | ?x AP61 cells<br>1x C6/36 cells |
|  |  |  | 2x Vero cells |
| DAK-AR-71 |  | *Aedes taylori* (Senegal 1984) | ?x AP61 cells<br>1x C6/36 cells<br>2x Vero cells |
| PRVABC59 | Asian | Human (Puerto Rico 2015) | 5x Vero cells |
| FLR |  | Human (Columbia 2015) | 3x C6/36 cells |
| H/PAN/2016/BEI-259634 |  | Human (Panama 2016) | 4x Vero cells |
| H/PAN/2015/CDC/259359 |  | Human (Panama 2015) | 4x Vero cells |

For each strain in the panel, two independent stocks were amplified on Vero cells to account for stock to stock variation, and the sensitivity of the viral stocks to pretreatment with IFNα-2a or IFNβ (1000 U/mL) in A549 cells was determined (Fig 2; S1 Table). Both African-lineage and Asian-lineage viruses were more potently inhibited by IFNβ pre-treatment than IFNα, with viral replication as measured by viral titers reduced 3 to 42-fold in response to IFNα and 63 to 807-fold in response to IFNβ (Fig 2A). The biggest differences were between DAK-AR-67 (African-linage) and FLR (Asian-lineage) (14-fold) for IFNα and DAK-AR-25 and MR 766 (both African-lineage, 13-fold) for IFNβ. There was a range of responses within each lineage: for example, among African-lineage strains, the most sensitive MR 766 isolate was more susceptible to both IFNα and IFNβ than another African lineage virus, IbH 30656 (3 and 12-fold respectively; Fig 2B). Similarly, among Asian-lineage strains, FLR was the most sensitive to IFNα and was 11-fold more susceptible than H/PAN/BEI-259634, while PRVABC59 and H/PAN/CDC-259359 were both equally susceptible to IFNβ and were 7-fold more sensitive than H/PAN/BEI-259634.

**Fig 2.**
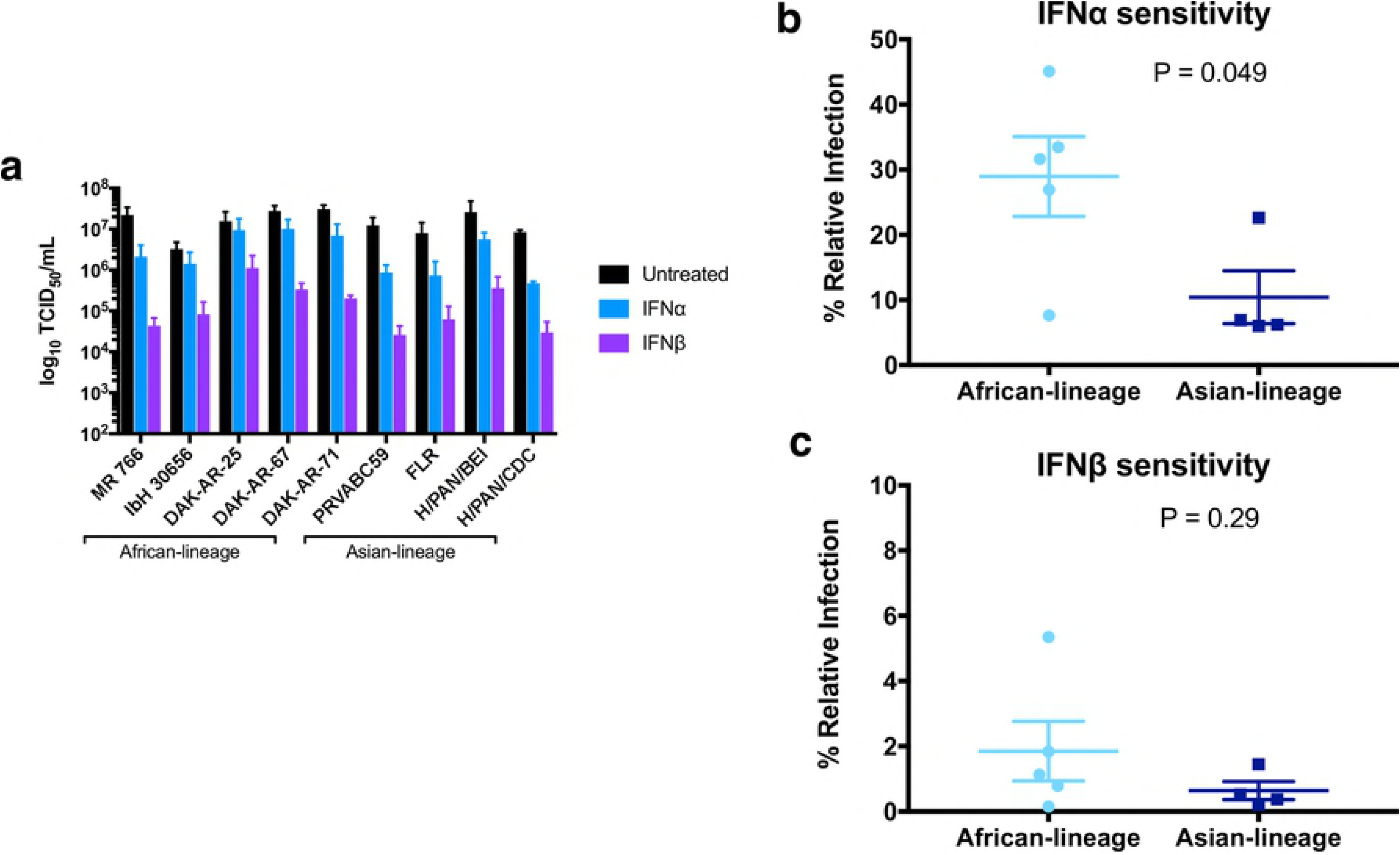
Effect of IFN-I pre-treatment on diverse Zika virus strains in A549 cells. (a) The susceptibility of each ZIKV strain to restriction by IFNα-2a or IFNβ was assessed in A549 cells. The titer (TCID50/mL) of each strain in the absence of IFN-I (black), presence of 1000 U/mL of IFNα-2a (blue), and presence of 1000 U/mL of IFNβ (purple) is shown. All data represent the average of four independent experiments that were carried out with two independently-generated stocks of each ZIKV strain. Error bars represent standard deviation. (b, c) Comparison of IFNα-2a-mediated (b) and IFNβ-mediated (c) restriction of African-lineage vs. Asian-lineage ZIKV strains. Percent relative infection (IFN+/IFN-) is plotted for African-lineage (light blue) and Asian-lineage (dark blue) ZIKV strains. Data points represent the average of four independent experiments that were carried out with two independently-generated stocks of each ZIKV strain. Error bars indicate the SEM. A two-tailed student’s t-test was used to compare percent relative infection of African-lineage vs. Asian-lineage ZIKV strains.

As an aggregate, African-lineage ZIKV strains were significantly less susceptible to IFNα restriction than Asian-lineage strains (Fig 2B; p=0.049); they were also less susceptible to IFNβ, though differences in sensitivity to IFNβ (Fig 2C; p=0.29) were not significant and were largely driven by one virus (DAK-AR-25) with high resistance. Taken together, the data reinforces IFN-I as a potent restrictor of ZIKV replication, albeit with substantial strain-to-strain differences in susceptibility, with African-lineage strains less sensitive to IFN-I than Asian-lineage strains.

### Expression of IFITM3 at levels similar to IFN-induction in A549 cells does not restrict ZIKV

Given the potent IFN-I-induced restriction of ZIKV strains in the panel, the role of the first ISG shown to restrict ZIKV was characterized. IFITM3 has been recently described as a potent IFN-I-inducible ZIKV restriction factor in HeLa, 293T, A549, HDFa, and HFF cells [28, 29]. IFITM3 is induced by both IFNβ and IFNα in A549 cells, with slightly higher levels (~2- fold) in IFNβ than IFNα -treated cells at the same dose (1000U/mL). The induction was dose dependent, as shown with increasing doses of IFNβ (Fig 3A). To determine whether the induction of IFITM3 expression could explain the sensitivity of ZIKV to IFN-I, an A549 cell line expressing an N-terminally FLAG-tagged IFITM3 was generated (Fig 3B). To ensure that the levels of IFITM3 were physiologically relevant, we sorted cells and selected cells with relatively lower levels of IFITM3. IFITM3-expressing A549 cells expressed similar (~ 2-fold higher) levels of IFITM3 than in IFNβ-treated A549 control cells (Fig 3C).

**Fig 3.**
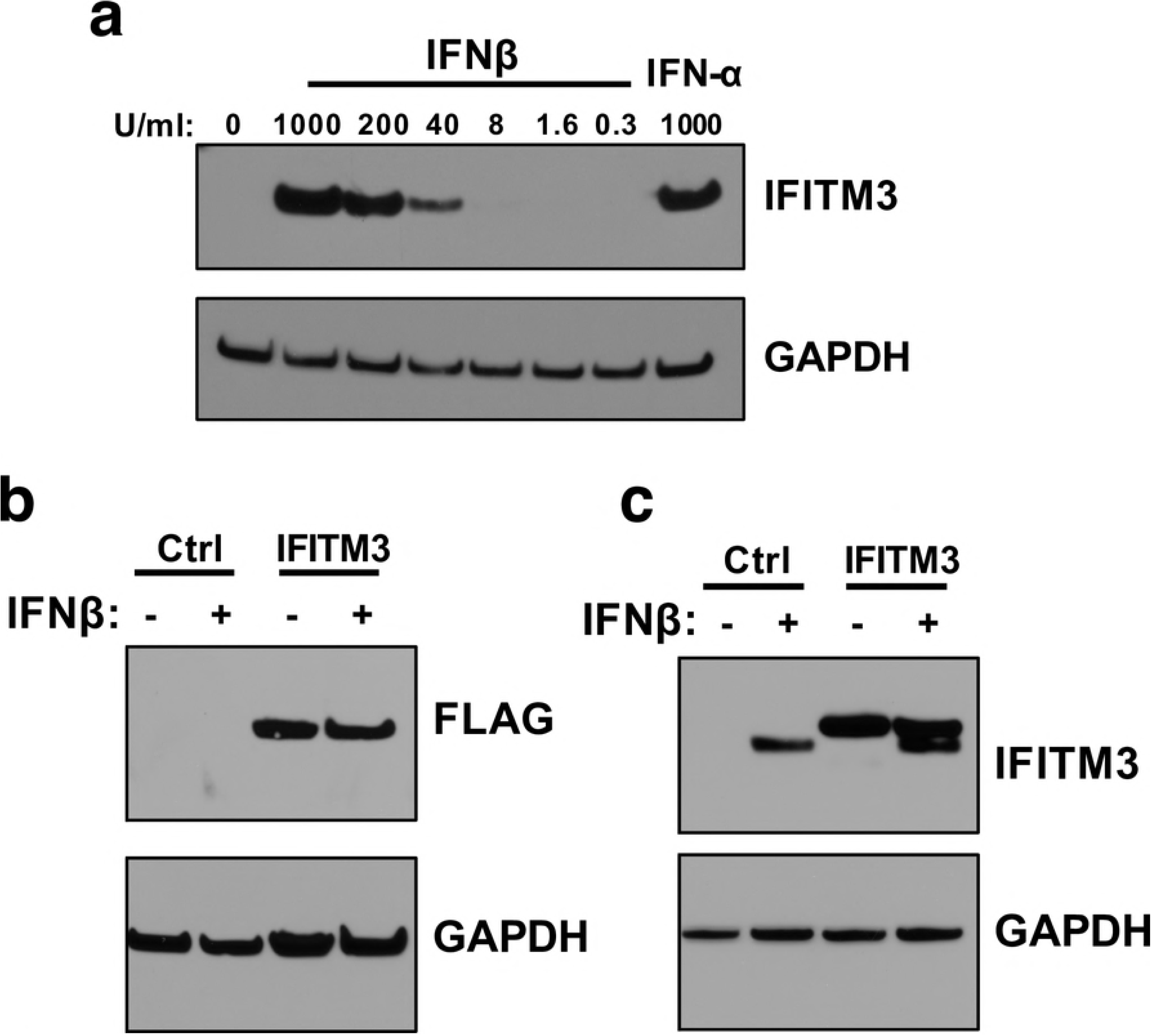
Expression of IFITM3 in A549 cells transduced with exogenous IFITM3 compared to after IFN-I-induction. (a) Western blot analysis of IFITM3 expression in A549 cells pretreated with increasing concentrations of IFNβ for 24 hours. The concentration of IFN is indicated above each lane. (b) Western blot analysis of IFITM3-FLAG expression using an anti-FLAG antibody in IFITM3 A549 cell lines. (c) Western blot analyses of expression of IFITM3-FLAG protein compared to endogenous IFITM3 using an anti-IFITM3 antibody.

To assess the effect of IFITM3 expression on ZIKV replication, IFITM3-expressing and control cells were infected with African-lineage isolate MR 766 and Asian-lineage isolate PRVABC59, both of which were found to be especially susceptible to IFN-I. Viral replication was not significantly different in cells expressing IFITM3 than from control cells for either strain (Fig 4A, B). Importantly, an Influenza A reporter virus was potently restricted in IFITM3-expressing cells, while virus-like particles expressing the murine leukemia virus envelope protein were not restricted in these cells (Fig 4C, D). This is consistent with published data showing IFITM3 restricts Influenza A virus but not murine leukemia virus [31].

**Fig 4.**
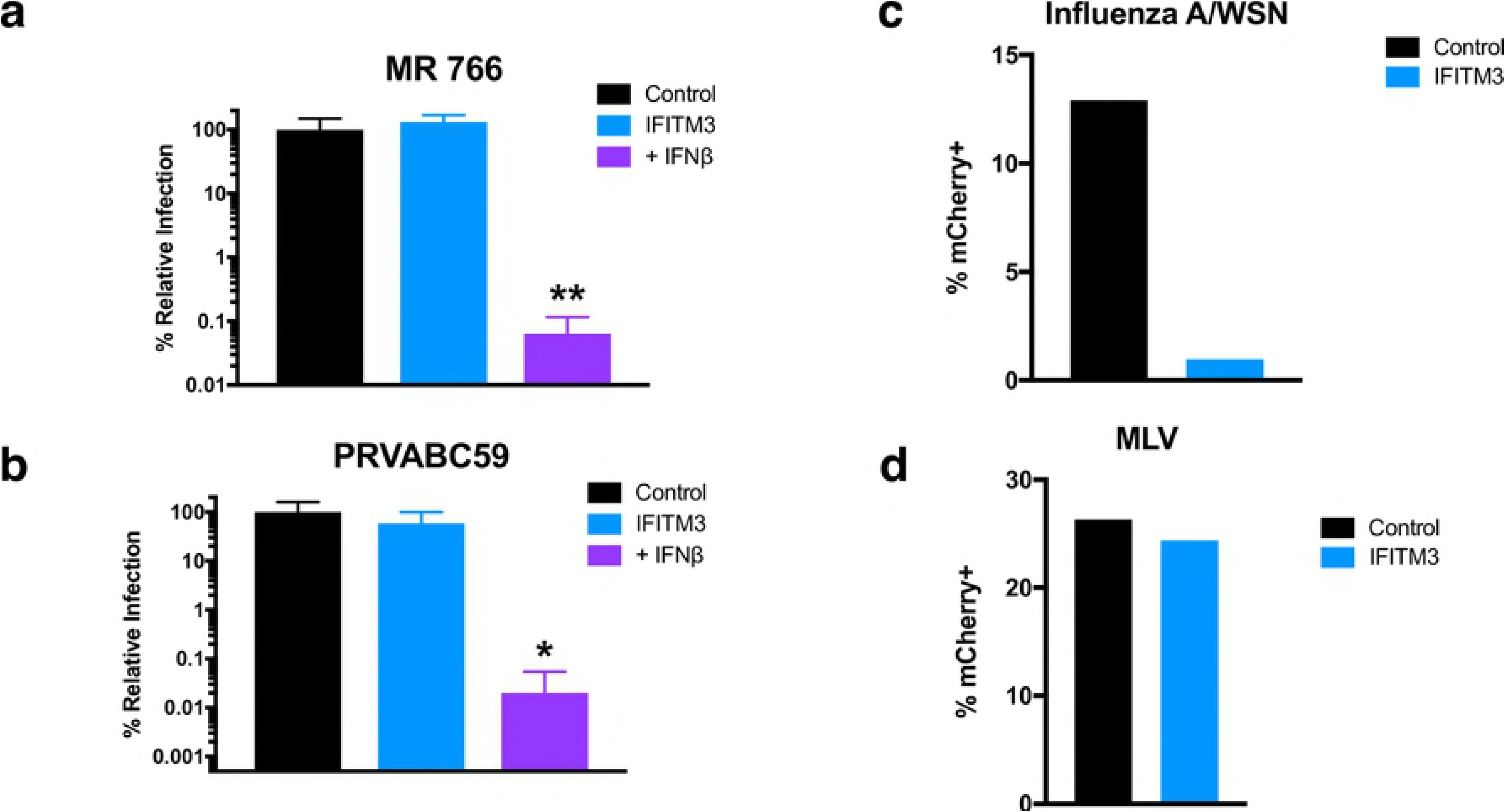
Infection of cells expressing IFITM3-FLAG in the absence and presence of IFNβ. (a, b) Infection with (a) MR 766 and (b) PRVABC59. Both panels show viral titers 48 hpi in untreated IFITM3 cells (blue) or pre-treated with 1000 U/mL of IFNβ (purple). For each strain, percent relative infection (IFITM3/Control or IFN+/Control) is shown. All data represent the average of four independent experiments that were carried out with two independently-generated stocks of each ZIKV strain. Error bars indicate standard deviation. (c, d) Infection of IFITM3 cells with mCherry-expressing (c) Influenza A/WSN reporter virus and (d) VLPs expressing murine leukemia virus envelope protein. *p < 0.05, **p < 0.01 (one-way analysis of variance (ANOVA) followed by Dunnett’s post-hoc test for multiple comparisons).

Notably, cells expressing IFITM3 continued to have a robust response to IFNβ, with drastic reductions in viral replication for both strains (1.6×10^3^ – 5.1×10^3^–fold) when compared to infection of untreated control cells (Fig 4A, B). Taken together, these data suggest that the physiological levels of IFITM3 induced by IFNβ in A549 cells are not sufficient to substantially antagonize ZIKV replication and may explain only a small part of the IFNβ effect in these cells.

### IFITM3 does not explain the IFN-I inhibition of ZIKV in A549 cells

ZIKV was not restricted in cells exogenously expressing IFITM3 at levels that appeared by Western blot to be similar to those induced by IFNβ, suggesting that IFITM3 may not be a major contributor to the IFNβ effect against Zika virus. However, we cannot rule out that the presence of a FLAG epitope impacts IFITM3 activity. Thus, to better define the contribution of IFITM3 to the overall IFN-I response against ZIKV replication, we employed a complementary CRISPR-Cas9 gene editing approach to knock out IFITM3. A549 cells were transduced with virus-like particles (VLPs) carrying a lentiviral vector encoding Cas9 along with IFITM3-targeting sgRNAs (sgRNA1 or sgRNA2) or a non-targeting control (NTC) sgRNA. Cells transduced with sgRNA1 or sgRNA2 were depleted in IFITM3 expression as compared to NTC cells, both basally and when treated with IFNβ (Fig 5A). Tracking of Indels by Decomposition (TIDE) analysis of cells transduced with IFITM3-targeting sgRNAs indicated that ~96% of sgRNA1-transduced cells and 88% of sgRNA2-transduced cells were edited at the IFITM3 locus (S1 Fig) [32]. Due to the high level of sequence identity between IFITM2 and IFITM3, IFITM2 expression was also knocked out in cells transduced with IFITM3-targeted sgRNAs (Fig S2). However, IFITM2 is not thought to impart ZIKV restriction [29], thus it was reasoned that the cell lines would be suitable to shed light specifically on the contribution of IFITM3 to the IFN-I response against ZIKV.

**Fig 5.**
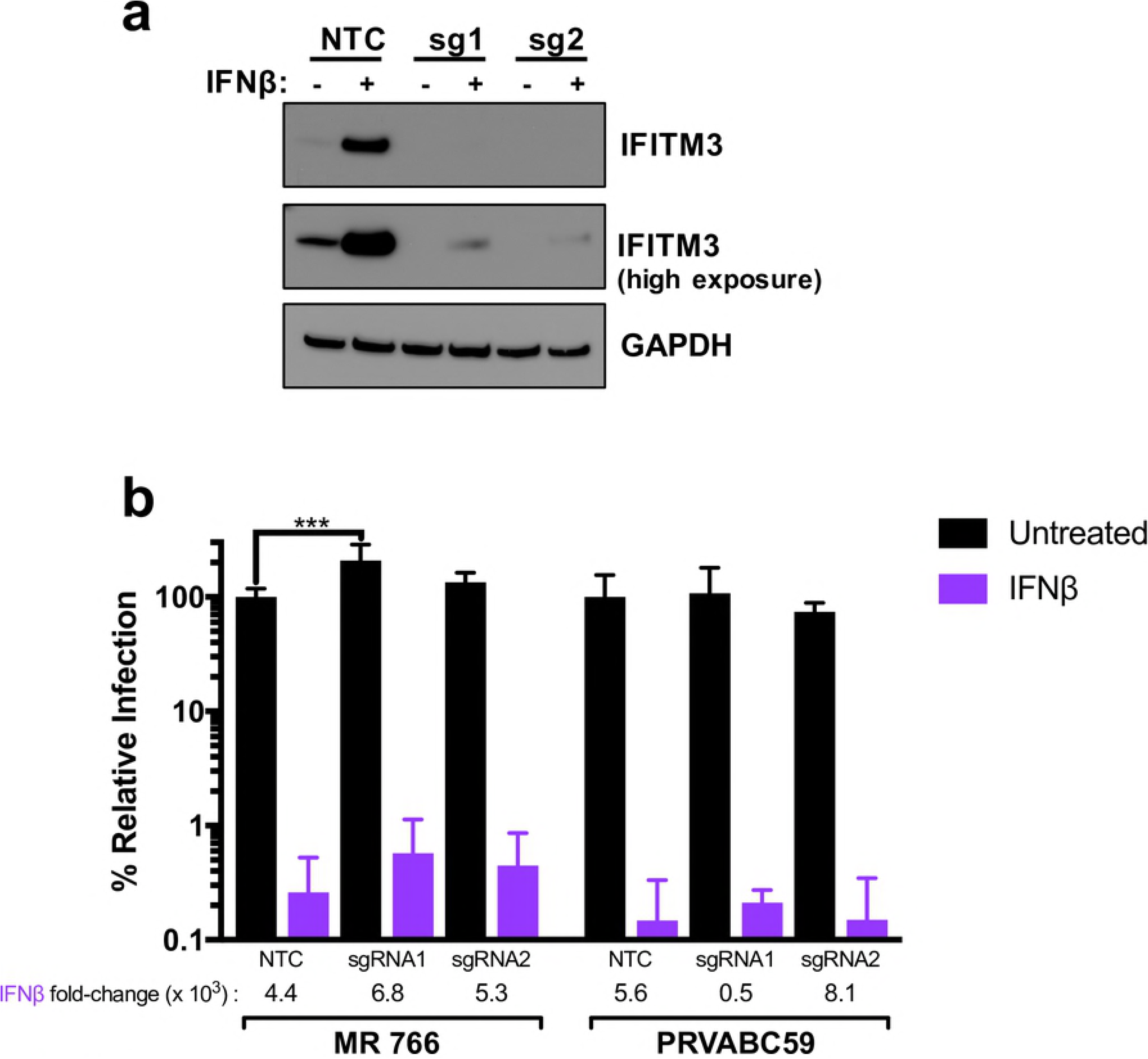
Analysis and infection results of IFITM 3 knock-out cells. (a) Western blot analysis of IFITM3 expression in untreated or IFNβ pretreated (1000 U/mL) A549 cells transduced with non-targeting control (NTC), sgRNA1 (sg1), or sgRNA2 (sg2). The sgRNA used to transduce each cell line is indicated above each lane. (b) Infection results with MR 766 and PRVABC59 showing viral titers 48 hpi in IFITM3-knockout and control A549 cells. The percent relative infection (normalized to NTC Untreated) of each strain in the absence of IFN-I (black) and presence of 1000 U/mL of IFNβ (purple) is shown in each indicated cell type. All data represent the average of four independent experiments that were carried out with two independently-generated stocks of each ZIKV strain. Error bars indicate standard deviation. ***p < 0.001 (two-way analysis of variance (ANOVA) followed by Dunnet’s post-hoc test for multiple comparisons).

To this end, IFITM3-knockout or NTC cells were infected with MR 766 or PRVABC59 at an MOI of 1. While there was a significant increase in viral replication of MR 766 in sgRNA1 IFITM3-knockout cells, the overall magnitude was small (~2.5-fold) and was not observed in sgRNA2 IFITM3-knockout cells infected with MR 766 (~1.4-fold). These slight increases in viral replication were also not consistently observed in IFITM3-knockout cells infected with PRVABC59. Moreover, when IFITM3-knockout cells were treated with IFNβ, there was a substantial decrease in viral replication as compared to untreated cells, with viral titers in IFITM3-knockout cells being decreased between 5×10^2^ and 8.1×10^3^-fold (Fig 5B). Viral replication was decreased by a similar amount (4.4×10^3^ and 5.6×10^3^-fold) in NTC cells treated with IFNβ and was not significantly different from IFITM3-knockout cells. This suggests at best a very modest contribution of IFITM3 to the overall IFN-I response to ZIKV in A549 cells, which is consistent with the results in cells exogenously expressing IFITM3, more significantly highlighting the major contribution of ISGs other than IFITM3, at least in A549 cells (Fig 5B).

## Discussion

We took advantage of a panel of African- and Asian-lineage ZIKV strains to define the IFN-I sensitivity of different ZIKV variants and determine if the host protein IFITM3 plays a critical role in the IFN-I response. ZIKV strains had a range of susceptibilities to IFN-I in A549 cells but both over-expression and knockout approaches suggest that IFITM3 does not play a major role in the IFN-I-induced restriction of ZIKV. African-lineage strains as a whole were more resistant to IFNα-mediated restriction when compared to Asian-lineage strains. This suggests a lineage-dependent phenotypic difference that is critical at the host-virus interface and supports future studies to identify ISGs that play important roles in restricting ZIKV replication.

The finding that African-lineage strains in the panel were more resistant to the effects of IFNα as compared to Asian-lineage strains was counter-intuitive given the fact that it is Asian-lineage strains that cause severe neuropathologic outcomes in fetuses, neonates, and adults. However, the data are in line with several recent studies that have demonstrated that infection with African-lineage strains results in enhanced replication kinetics, virus production, cytopathicity, and disease progression in murine models [10-22]. Further, one of these studies has shown that induction of IFN-I is higher following infection with African-lineage strains in murine models [20]. While IFN is a potent antiviral protein, it also plays an important role in immune activation and for this reason can have dual roles in viral infection outcomes. One hypothesis to explain these data is that decreased virulence, reduced immune activation, and IFN-I sensitivity may be conducive to establishing persistent infections within certain tissues and that rapid, self-limiting virus replication may minimize opportunities to establish infected cell sanctuaries. Indeed, others have suggested that Asian-lineage strains may be better able to establish chronic infection of neural progenitor cells, undergo more efficient vertical transmission, and establish viral reservoirs in the central nervous system, lymph nodes, and gastrointestinal and genitourinary tracts [22, 33-36]. A caveat of the current study is the divergent passage histories of the strains tested. Future studies with larger numbers of low-passage strains involving important target cell types of ZIKV tropism and pathogenesis will be critical in strengthening our understanding of these relationships.

All ZIKV strains were potently restricted by IFNβ pre-treatment. Although IFNα and IFNβ signal through the same heterodimeric IFN receptor (IFNAR), IFNβ has been reported to possess higher binding affinities for IFNAR and can independently bind one IFNAR chain (IFNAR1), which triggers the downstream expression of a unique set of ISGs [37, 38]. A combination of these factors likely influence the stronger potency of IFNβ-mediated viral restriction we and others have observed [39]. Of note, we tested two (IFNα-2a and IFNβ) of the many subtypes of IFN-I that have been shown to have specific and distinct biological effects [38, 40-42]. It will be important to test other IFN-Is alone and in combination in future studies of IFN-I-induced restriction of ZIKV.

It is noteworthy that while the African-lineage strains were overall more resistant to IFNα than the Asian-lineage strains, one commonly used African isolate, MR 766, was very sensitive to both IFNα (19-fold knockdown) and IFNβ (807-fold knockdown). This may reflect the extensive passage history of MR 766, which could have selected for a virus that is, as a result, not adapted to evade innate immune pressures. Of note, the differences between African-lineage and Asian-lineage groups in terms of their IFNα would have been even stronger if MR 766 were not included in our panel. In addition, it is interesting that two of the African-lineage isolates tested, MR 766 and DAK-AR-25, are not very divergent (Fig 1) with only 38 amino-acid differences between the two isolates throughout their entire coding sequences, yet DAK-AR-25 was one of the most IFN-I-resistant viruses (IFNα=5-fold; IFNβ=63-fold). Recently, key sequence determinants in NS1, for increased evasion of IFN-I signaling, and NS4B, for replication in immunocompetent mice, have been identified [43, 44]. However, these determinants do not account for the differences between MR766 and DAK-AR-25 observed in this study. Thus, comparative sequence analysis of the panel strains and individual stocks could be leveraged to identify key sequence determinants that predict relative sensitivity to IFN-I.

IFITM3 has recently been reported as an important ZIKV-restricting host factor that blocks an early stage of the ZIKV replication cycle [28, 29]. The current findings provide evidence that IFITM3 does not play a substantial role in the IFN-I-induced restriction of ZIKV, at least in A549 cells where there is potent IFN-induced inhibition of ZIKV. Two ZIKV isolates belonging to each lineage (MR 766 and PRVABC59), that were highly sensitive to IFN-I, were not significantly restricted by IFITM3 when it was expressed at physiologically-relevant levels. Two viruses previously shown to be either IFITM3-susceptible (IAV) or resistant (MLV) behaved as expected when they were used to infected IFITM3-expressing cells [31]. Pre-treatment of the IFITM3-expressing cell line with IFNβ resulted in potent restriction of ZIKV strains, underscoring the critical contribution of ISGs other than IFITM3 in the IFN-I-mediated restriction of ZIKV. ZIKV restriction also remained unchanged in IFNβ-treated cells in which endogenous IFITM3 had been knocked out, providing further support that endogenous levels of IFITM3 induced by IFN-I do not play a critical role in restricting ZIKV replication. Of note, we observed a 1.4 – 2.5-fold increase in viral replication in IFITM3-deficient cells infected with MR 766, which is consistent with previous studies that reported an increase in percent of infected cells after shRNA-mediated knockdown of IFITM3 [28] or deletion of the *Ifitm3* locus in murine embryonic fibroblasts [29]. Knocking out IFITM3 expression also abrogated IFITM2 expression. Thus, because there were not significant differences in viral replication between IFITM2/3-deficient cells and control cells, the data is consistent with others who have found that IFITM2 does not play an important role in the IFN-I-induced restriction of ZIKV [28].

There are several reasons that may explain why the current findings differ from previous studies that have described an important role for IFITM3 in restricting ZIKV. First, we utilized different methods for quantifying ZIKV replication. We measured viral titers by TCID_50_ assay in cell supernatants, while previous reports have used ZIKV envelope glycoprotein or double-stranded DNA immunofluorescence. These assays capture different aspects of ZIKV replication and could explain some of our divergent findings although based on the proposed mechanism of action of IFITM3 early in the life cycle, this would not be expected to be a major factor. Importantly, we used an MOI of 1 and measured viral replication at 48 hpi, which is consistent with most experiments performed in these previous studies. However, prior studies have used HeLa and MEF cells for some experiments to describe IFITM3-mediated restriction of ZIKV. Others have found that IFITM3-mediated restriction can depend on cell type, raising the interesting possibility that IFITM3 restricts ZIKV infection in some cells but not others [30]. A potentially critical difference that may explain the differences in results is the fact that several of these studies have utilized systems that, in many cases, highly overexpress IFITM3 to levels much higher than seen upon IFN-I induction [28, 29]. Thus, it is quite possible that IFITM3 can restrict infection when present at very high levels, although this may not recapitulate the response to IFN-I, where IFITM3 is not expressed at such high levels. It is also worth noting that ours is the first study to examine IFITM3-mediated restriction of ZIKV using a complete Cas9-mediated knockout as opposed to a shRNA/siRNA-mediated knockdown approach, including a knock-down that targeted the overexpressed IFITM3 gene in the same cell line [28, 29].

Overall, the results of this study demonstrate that the effects of IFNα on ZIKV replication in A549 cells are lineage-dependent in our panel of ZIKV strains. The inter-strain variation in IFN-I sensitivity across all viruses in the panel is intriguing and future studies using these strains may identify determinants of IFN-I sensitivity and/or resistance. Finally, the findings indicate that IFITM3 does not play a significant role in the IFN-I-induced restriction of ZIKV replication and thus support continued investigation of ISGs that do restrict ZIKV. Recent studies using targeted approaches against known restriction factors have reported the ability of two IFN-I-inducible host anti-viral proteins to restrict ZIKV [45-47]. In addition to targeted screens focused on known anti-viral host proteins, large-scale loss or gain-of-function genetic screens will help to identify these as-yet unknown innate immune factors.

## Materials and methods

### Viruses

Zika virus strains were kindly provided by BEI Resources (MR 766, IbH 30656, PRVABC59, FLR, H/PAN/2016/BEI-259634, H/PAN/2015/CDC-259359) and Michael Diamond (DAK-AR-25, DAK-AR-67, DAK-AR-71). All strains were tested for mycoplasma using the Universal Mycoplasma Detection Kit (ATCC) by inoculating 1e6 HEK293T cells with 100 uL of viral aliquot and harvesting cells and cell debris 5 days post-inoculation according to the manufacturer’s protocol. Any stocks found to have mycoplasma contamination were filtered through a 0.2 um filter and re-tested and confirmed to be mycoplasma-free before proceeding with virus propagation. All Zika virus strains were propagated by inoculating Vero cells at an MOI of 0.01 in a minimal volume of serum-free DMEM for 4-6 hours. After inoculation, fresh DMEM supplemented with 2mM L-glutamine, 1X Anti-anti (anti-microbial/anti-mycotic, Gibco) and 3.3% FBS was added to the inoculum so that the final concentration of FBS was 2%. Supernatants were collected 4 days post-inoculation, except in the case of the FLR isolate, which gave higher titers when supernatants were collected 5 days post-inoculation. Supernatants were cleared of cellular debris by centrifuging for 10 minutes 300g at 4°C, before aliquoting and storage at −80°C and viral titers were determined by the TCID_50_ assay described below. Experiments were performed with aliquots that had undergone at most two freeze-thaw cycles, which was not found to have any discernible effect on viral titers.

### Cells

A549 cells (A. Berger; ATCC) were maintained in RPMI (Invitrogren) supplemented with 10% fetal bovine serum (FBS), 2mM L-glutamine, and 1X Anti-anti (anti-microbial/anti-mycotic, Gibco). Vero cells (A. Geballe; ATCC) and HEK293T cells were maintained in DMEM (Invitrogen) supplemented with 10% FBS,2mM L-glutamine, and 1X Anti-anti. The identity of the A549 cells and HEK293T cells was confirmed using STR CODIS finger-printing and all cell lines were found to be mycoplasma-free by the Research Cell Bank shared resource at the Fred Hutchinson Cancer Research Center.

### Sequencing and phylogenetic analysis of ZIKV strains

All Zika virus stocks were sequence-confirmed by Sanger sequencing of a 1.8 – 3.4-kbp region of the Zika virus genome that encodes non-structural proteins 1 through 3. The complete open reading frame of DAK-AR-67 was sequenced, since there was no sequence data available for this isolate. To do this, viral RNA was isolated using the QiaAMP Viral RNA Mini Kit (Qiagen) and cDNA was produced using SuperScript III First Strand Synthesis System (Invitrogen) with random hexamers according to the manufacturer’s suggested protocol. The Primal Scheme primer designer software <u>(http://primal.zibraproject.org/)</u> was then used to design primers that tiled across the complete open reading frame in ~645 bp fragments that overlapped by ~210 bp (Table S2) [48]. Five overlapping amplicons were generated by PCR amplification of cDNA with Q5 ReadyMix (NEB) using a subset of primer pairs (Table S3). Thermocycling conditions used were:

98°C, 30 s
98°C, 15 s, 30x
65°C, 5 min.

Each of the amplicons was then subjected to Sanger sequencing using the primers indicated in Table S2. Full-length open-reading-frame nucleotide sequences of ZIKV strains in the panel, as well as other ZIKV strains, were used to construct a maximum-likelihood phylogenetic tree with PhyML using a general time-reversible nucleotide substation model [49].

### IFN-I sensitivity assay

For each ZIKV strain, 8×10^4^ A549 cells were plated in each of three wells of a 24-well plate in a final volume of 1 mL of complete RPMI. One well was left untreated and the other two wells were pretreated with 1000 U/mL of IFNα-2a or IFNβ for 24 hours. After pre-treatment, cells were infected at an MOI of 1 in a final volume of 250 µL of serum-free RPMI for 4-6 hours. The inoculum was then aspirated, cells were washed twice with 1X PBS, and replenished with 1 mL of complete RPMI without IFN-I or containing 1000 U/mL of IFNα-2a or IFNβ. At 48 hours post-infection (hpi), 250 uL of supernatants were harvested and cleared of cellular debris at 4°C at 300G for 10 minutes and 2 × 100 uL aliquots were stored at −80°C until titration by TCID_50_ assay. All infections were performed with two separately-generated stocks of each ZIKV strain with biological duplicates for each stock. For the data analysis, all values were plotted and statistical analyses performed using Prism version 7 (GraphPad Software). TCID_50_/mL and Percent Relative Infection were calculated. Percent Relative Infection was determined by dividing the titer in the IFNα- or IFNβ-treated sample by the untreated sample.

### TCID_50_ assays

Zika viral titers were determined by TCID_50_ assay on Vero cells in a 96-well format. One day prior to titration, Vero cells were seeded in 100 uL of complete DMEM in a flat-bottomed 96-well plate at 8×10^3^ cells per well. For each condition tested, seven serial 10-fold dilutions of viral supernatants were prepared, starting at a concentration of 1 uL/well, with each dilution including 10 replicate wells and 2 mock infected wells. Cells were infected with 50 uL of each viral dilution in serum free DMEM for 4-6 hours, before being replenished with 100 uL of DMEM with 3% FBS, for a final concentration of 2% FBS. On day 5 post-infection the wells at a given dilution were scored by light microscope for the presence or absence of cytotoxicity and the TCID_50_/mL was calculated using the Spearman-Karber method.

### Generation of stable cell lines overexpressing IFITM3

IFITM3-expressing A549 cells were generated as previously described [50]. Briefly, the N-terminal FLAG-tagged IFITM3 open-reading frame was cloned into pHIV-Zsgreen directly upstream of the IRES-driven ZsGreen fluorescent reporter. Virus-like particles (VLPs) were generated in HEK293T cells by co-transfecting cells with pHIV-ZsGreen constructs (either IFITM3-encoding or empty vector as control) [51], psPAX2 (HIV-based packaging plasmid) [52], and pMD.G (vesicular stomatitis virus glycoprotein [VSV-G] envelope plasmid) [53] at a ratio of 1:1:0.5 using FuGENE 6 (Promega) according to the manufacturer’s protocol. Supernatants from HEK293T cells were collected 48 hours post-transfection and concentrated ~100-fold using Amicon Ultracel 100 K filters (Millipore). VLPs were then used to transduce A549 cells that has been plated 24 hours prior in a 6-well plate at 1×10^5^ cells/well in 2 mL of RPMI supplemented with 10% FBS and 2mM glutamine. A549 cells were transduced by spinoculation at 1200 × g for 90 minutes. The following day, the cells were expanded into new T75 flasks and were subsequently passaged and maintained in complete DMEM. IFITM3-expressing cells were sorted by gating cells in the fiftieth-percentile of zsGreen expression on a FACSAria II cell sorter.

### Generation of IFITM3 knockout cells lines

For generation of IFITM3-knockout A549 cell lines, guide RNAs targeting the first exon of *Ifitm3*, or non-targeting control guide RNA, were cloned into pLentiCRISPR, which allows for the expression of the guide RNA and Cas9 from the same construct. VLPs were generated as described above by co-transfecting the pLentiCRISPR plasmids, the psPAX2 packaging vector, and pMD2.G (vesicular stomatitis virus glycoprotein [VSV-G] envelope plasmid) at a ratio of 1:1:0.5 using the FuGENE 6 transfection reagent (Promega) according to the manufacturer’s protocol. The following day, cells were expanded into new T75 flasks and cultured. Subsequently, cells were passaged and cultured in complete media supplemented with 2 µg/ml of puromycin to select for sgRNA and Cas9 expression. The two sgRNAs that yielded the most efficient knockout of IFITM3 were sgRNA1, 5’-GCAGCAGGGGTTCATGAAGA-3’; and sgRNA2, 5’-TTGAGCATCTCATAGTTGGG-3’ and the non-targeting control was 5’-ATCTCGGGTCGACTGCGGAT-3’. Gene knockout was characterized by Tracking of Indels by Decomposition (TIDE) analysis. Briefly, after three rounds of puromycin selection, genomic DNA was isolated using the QuickExtract DNA extraction solution (Lucigen) by resuspending cells in 100 μL of the solution, and by denaturing for 20 min at 60 °C and 20 min at 95 °C. The *ifitm3* locus was amplified using the following primer set: forward ACCATCCCAGTAACCCGACCG and reverse GCTGATACAGGACTCGGCTCC. Amplicons were Sanger sequenced and gene editing was measured using TIDE analysis (https://tide-calculator.nki.nl/).

### Western blots and quantification

Whole cell extracts were prepared by lysing the cells in RIPA cell lysis buffer (50 mM Tris pH 8.0, 0.1% SDS, 1% Triton-X, 150 mM NaCl, 1% deoxycholic acid, 2 mM PMSF). Standard Western blotting procedures were used with the following antibodies: IFITM3 (Proteintech 11714-1-AP, used at 1:1000 dilution), IFITM2 (Proteintech 66137-1-Ig, used at 1:500 dilution), FLAG (OriGene TA100023, used at 1:2000 to 1:5000 dilution), and GAPDH (BioRad MCA4739P, used at 1:5000 dilution). Protein expression was quantified by measuring the band intensities using LI-COR Image Studio Software.

### Influenza A virus and Murine Leukemia Virus VLP infections

Influenza A virus/WSN (IAV; generously provided by A. Russell and J. Bloom) is an mCherry-expressing reporter virus where HA is replaced with mCherry. For murine leukemia virus (MLV), reporter VLPs were made by packaging the lentiGuide.mCherry vector [54] (a gift from Richard Young, AddGene plasmid #104375) with psPAX2 and pseudotyping with an amphotropic MLV envelope. For both viruses, 8×10^4^ IFITM3-expressing and control cells were plated in a 24-well plate one day prior to infections in a final volume of 1 mL of complete RPMI. For IAV, cells were infected at an MOI of 10 in 500 µL of complete RPMI for 16 hours. Cells were harvested and fixed in 1% paraformaldehyde. For MLV, cells were infected with a dilution of VLPs in complete RPMI supplemented with 10 µg/mL DEAE dextran. Cells were harvested and fixed in 1% paraformaldehyde 72 hours post infection. Both IAV and MLV-infected cells were assessed for mCherry expression using a Fortessa X50 flow cytometer and data was analyzed using FlowJo v9 software.

## Acknowledgments

We thank Michael Diamond for providing DAK-AR-25, DAR-AR-67, and DAK-AR-71 Zika virus strains; Alice Berger for providing the A549 cell line; Adam Geballe for providing Vero cells; Alistair Russell and Jesse Bloom for providing the Influenza/WSN-mCherry reporter virus; Adam Geballe, Nicholas Chesarino, and Michael Emerman for helpful discussions; Nicholas Chesarino for comments on the manuscript; and the NIAID’s Biodefense and Emerging Infectious Disease Resource Repository (BEI Resources) for providing MR 766 (WRCEVA), IbH 30656 (WRCEVA), PRVABC59 (BJ Russell), FLR (R Rico-Hesse), H/PAN/2016/BEI-259634 (BEI Resources), and H/PAN/2015/CDC-259359 (AM Powers).

## Supporting information

**S1 Table. Fold reduction in viral replication of each ZIKV strain by IFN-I.** The table lists the fold-reduction in viral replication and standard deviation (SD) of each ZIKV strain by pre-treatment of IFNα and IFNβ (1000 U/mL) in A549 cells.

**S2 Table. Zika virus sequencing primers**. The table lists all primers used to sequence Zika virus strains included in this study.

**S3 Table. Zika virus sub-amplicon primer pairs**. The table lists the primers pairs used to generate sub-amplicons for sequencing.

**S1 Fig. Tracking of Indels by Decomposition analysis of IFITM3-knockout A549 cells.** (a, b) TIDE analysis of sgRNA1-transduced (a) and sgRNA2-transduced (b) A549 cells. Deletion or insertion of nucleotides is plotted on the x-axis and percent of sequences is plotted on the y-axis.

**S2 Fig. IFITM2 expression is knocked out in IFITM3-knockout A549 cells.** Western blot analysis of IFITM2 expression in untreated or IFNβ pretreated (1000 U/mL) A549 cells transduced with non-targeting control (NTC), sgRNA1 (sg1), or sgRNA2 (sg2). The sgRNA used to transduce each cell line is indicated above each lane.

